# A Graph Matching Approach to Tracking Neurons in Freely-Moving *C. elegans*

**DOI:** 10.1101/2023.11.30.569341

**Authors:** Corinne Jones, Mahsa Barzegar-Keshteli, Alice Gross, Guillaume Obozinski, Sahand Jamal Rahi

**Affiliations:** Swiss Data Science Center, École polytechnique fédérale de Lausanne, 1015 Lausanne, Switzerland; Laboratory of the Physics of Biological Systems, École polytechnique fédérale de Lausanne, 1015 Lausanne, Switzerland

## Abstract

**Motivation:** Recent advances in 3D microscopy allow for recording the neurons in freely-moving *C. elegans* at high frame rates. In order to read out calcium activity, it is necessary to track individual neurons from frame to frame. However, doing this by hand for tens of neurons in a single ten-minute recording requires more than a hundred hours. Moreover, most methods proposed in the literature for tracking neurons focus on immobilized or partially-immobilized worms and fail with freely-behaving worms.

**Results:** In this paper we present an approach based on graph matching for tracking fluorescently-marked neurons in freely-moving *C. elegans*. Neurites (and sometimes neurons) can be oversegmented into pieces at the preprocessing phase; our algorithm allows several segments to match the same reference neuron or neurite. We demonstrate our method on three recordings. We find that with five labeled frames we can typically track the neurons and pieces of neurites with over 75% accuracy, with more reliable annotations for the most distinctive neurons.

**Availability and Implementation:** The code and preprocessed data will be made available upon publication.

## 1 Introduction

3D microscopy techniques allow researchers to record the calcium activity of many neurons of organisms such as *C. elegans* simultaneously at a high temporal resolution (Kato et al., 2015, Nguyen et al., 2016, Venkatachalam et al., 2016). The resultant videos are a rich source of information, and can potentially provide insight into computations of neural circuits. However, the current bottleneck lies in processing the videos: In order to read out the activity of the neurons, they must be tracked from frame to frame. *C. elegans* hermaphrodites have 302 neurons and a largely stereotyped brain (White et al., 1986, Zhen and Samuel, 2015, Witvliet et al., 2021) However, there are still differences from worm to worm, and differences also appear across time in videos of the same worm (Toyoshima et al., 2020, Yemini et al., 2021). It is important to study freely-moving worms, as immobilized worms behave differently (Hallinen et al., 2021). Moreover, it is also important to track their neurites, as meaningful information lies in the neurites of the worms, in addition to the nuclei and somas (Park et al., 2022).

### Contributions

In this work we focus on tracking somas and neurites in videos of freely-moving young adult *C. elegans*. Overall, we make four main contributions to the worm tracking literature. First, our paper is among the first to track freely-moving worms (with exceptions including the works of Yu et al. 2021 and Park et al. 2022). Such worms often deform, and without using an additional camera to help straighten the worm, the tracker must be able to directly handle such deformations. Moreover, the neurons can move in and out of the field of view. We overcome these hurdles by tracking the neurons using graph alignment techniques. In our approach, each frame is represented by a complete graph, with the somas and (parts of) neurites being the nodes. Our algorithm learns to use the node features (locations and shapes of the objects) and edge features (distances between the objects) to match the nodes in each frame with nodes in a reference frame. By using only relative position information, this approach is able to handle non-rigid deformations. Moreover, by relying on such features, the method provides interpretable results. Second, our method can recognize when multiple disjoint parts comprise a single object and is therefore able to track neurites. Our algorithm achieves this thanks to a novel formulation allowing for many-to-one matching. With the exception of the method of Park et al. (2022), existing approaches focus on tracking only neurons and assume that each neuron appears as a single object. Third, our method requires few labeled frames. Our algorithm uses the labeled frames in order to tune the hyperparameters. Deep learning approaches such as that of Park et al. (2022) generally rely on significantly more labeled frames and require longer training times. Finally, we propose a method for learning the relative importance of each of the different features used in order to optimize the quality of the tracking. We do this by leveraging techniques from structured output learning. Prior graph-based approaches left these values fixed (Kainmueller et al., 2014, Chaudhary et al., 2021, Toyoshima et al., 2020).

### Related work

There are three main steps in tracking: preprocessing the frames, segmenting the objects in each frame, and identifying the objects in each frame. Like most of the literature, we will take the preprocessing steps and segmentations as given and focus on the identification step. Three main strategies have been used to track *C. elegans* neurons in the past: statistical models (e.g., Nguyen et al., 2017, Varol et al., 2020, Nejatbakhsh and Varol, 2021, Wen et al., 2021, Tokunaga et al., 2014, Hirose et al., 2018, Chaudhary et al., 2021), graph-based approaches (e.g., Sutton and McCallum, 2012, Lyzinski et al., 2016, Swoboda et al., 2017, 2019, Kainmueller et al., 2014, Toyoshima et al., 2020, Chaudhary et al., 2021), and deep learning (e.g., Yu et al., 2021, Park et al., 2022); see Appendix A for more details. The method we will propose in the present paper is most similar to the methods of Kainmueller et al. (2014) and Chaudhary et al. (2021). While we also employ graph matching based on pairwise node and edge features, our paper differs in terms of the parts of the worms tracked and their movement, the features generated, the formulation, and the optimization. This is because we wish to analyze somas and neurites in freely-moving young adult worms. In contrast, Kainmueller et al. (2014) focus on the nuclei of pre-straightened worms at the L1 larva stage and Chaudhary et al. (2021) focus on nuclei of partially-immobilized worms. Both sets of authors either work with pre-straightened worms or need some manual intervention in order to align them. The features they use rely fundamentally on the worms being straightened, whereas ours do not require straightened worms and are rotation-invariant in the xy-plane. (Significant changes in the z-coordinates occurred rarely in our recordings.) Videos of freely-moving worms have additional complexities, for example, due to smearing and parts of the worms moving in and out of the frame. Moreover, (parts of) neurites can look much more distinct than nuclei. Due to this added complexity, the second main difference is that we propose additional node features that help distinguish between object parts.

There are also large differences in the formulation and optimization. As previously noted, multiple disjoint parts (e.g., pieces of a neurite) in an image may in reality form a single object (a neurite). Therefore, we explicitly add constraints that allow multiple objects in one frame to be matched to a single object in another frame. In terms of optimization, Kainmueller et al. (2014) apply the dual decomposition approach of Torresani et al. (2013) for one-to-one matching. In our case one would have to modify the dual decomposition approach for the many-to-one matching case, which would require developing a different rounding method for obtaining a final solution. In contrast, to address the scalability of the problem, we use a simple multi-step strategy and solve linear programming relaxations of the objective function for multiple sets of object sizes. In other scenarios, e.g., when all objects have a similar size, a modified version of the dual decomposition approach could be more suitable because it addresses scalability in a different way. In contrast, Chaudhary et al. (2021) used max-product loopy belief propagation. Torresani et al. (2013) found that for images from other domains, their method performed similarly to or better than max-product loopy belief propagation in terms of accuracy, but that it is slower in terms of runtime. However, in our case we require many-to-one matching constraints, and it is non-trivial to generalize loopy belief propagation to this setting, and to construct algorithms with convergence guarantees. We also demonstrate that one can successfully optimize the hyperparameters in graph matching to improve the performance when there are labeled frames present. While Chaudhary et al. (2021) mention this as a possibility for hyperparameters appearing linearly in the cost function, they do not try it in practice. In contrast, we note that it can be done even for parameters that modify the features non-linearly, and demonstrate in experiments that it can be beneficial. Thus, while graph matching has been used for tracking *C. elegans* neurons in the past, the existing methods do not allow for tracking freely-moving worms with objects that may sometimes appear as multiple disjoint parts. In addition, we propose a simple yet efficient way to make the algorithm scalable in this setting and to learn an optimal weighting of the different features.

## 2 Method

The three key components of our method are the features we use, the formulation of the graph matching problem, and the optimization approach for solving the graph matching problem. In this section we discuss each component in turn.

Our inputs are videos that have been preprocessed such that the objects in each frame are segmented. See Appendix C for how we performed the preprocessing on the videos used for this paper. In the present setting, the objects will consist of both somas and (parts of) neurites. Each soma and neurite may potentially be represented by multiple disconnected parts, and these parts are not a priori known to belong together. In the bottom left panel of Figure 2, the dark purple neurite illustrates this scenario: in the image the neurite is split into three disconnected parts, whereas we know that all three parts comprise the same neurite.

In addition, there is one frame, which we refer to as a reference frame, that contains all objects. In this frame the objects have been automatically segmented and then manually labeled such that multiple parts that all belong to the same object are given the same identity. We will align every frame to this reference frame.^1^ Aligning each frame to a reference frame avoids the problem of accumulating errors if one were to instead match each frame to the next frame. Moreover, having all objects exist in the reference frame means that one can perform many-to-one matching, matching anywhere from zero to several objects in a given frame to one object in the reference frame. Tackling this problem is significantly more straightforward than performing many-to-many matching and/or handling the case where neurons may be missing in both frames.

### 2.1 Features for graph matching

To match a given frame to the reference frame, we develop features related to the compatibility of nodes between the two frames and the compatibility of edges between the two frames. In order to be able to handle moving worms, we design features that are largely translation- and rotation-invariant.

Concretely, consider a node indexed by *i* in the given frame and a (potentially different) node indexed by *i*′ in the reference frame. Define the eccentricity of an object to be the eccentricity of the best-fitting ellipse to the projection of the object on the x-y plane. Furthermore, define the solidity of an object to be the ratio of its volume to the volume of its convex hull (Russ, 2011, Chapter 11). We propose to generate the node similarities *K*_*v*_(*i, i*′) of every potential match (*i, i*′) by comparing the centers of mass *z*_*i*_, *z*_*i*_′ in the z-direction (which tend to be stable across frames), the eccentricities *ε*_*i*_, *ε*_*i*_′ and the solidities *s*_*i*_, *s*_*i*_′ :

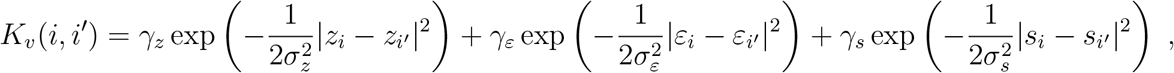

where *γ*_*z*_, *γ*_*ε*_, and *γ*_*s*_ are weighting hyperparameters and *σ*_*z*_, *σ*_*ε*_ and *σ*_*s*_ are bandwidth hyperparameters that need to be set. If multiple labeled frames are available, all of these hyperparameters can be learned; see Section 2.5. Otherwise, we can assign each term equal weight and set the bandwidths using the median pairwise distance between the inputs, a common rule of thumb (Fukumizu et al., 2009).

We define the compatibility between edges based on the distances between nodes. Concretely, let *i* and *j* be two nodes in the given frame and *i*′ and *j*′ be two nodes in the reference frame. Denote the lengths of the edges from nodes *i* to *j* and from nodes *i*′ to *j*′ by *e*_*ij*_ and *e*_*i*_′*j*′, respectively. Then we define the edge compatibility matrix by

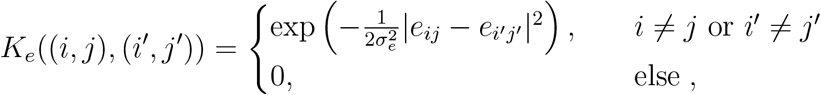

where *σ*_*e*_ is a bandwidth hyperparameter that needs to be set. Once again, if there are multiple labeled frames available, *σ*_*e*_ can be learned. Otherwise, it can be set using the median pairwise distance heuristic. Note that the case where *i* = *j* and *i*′ = *j*′ corresponds to comparing the compatibilities between nodes (since *e*_*ij*_ = *e*_*i*_′*j*′ = 0), which is covered by the matrix *K*_*v*_. See Figure 1 for an illustration of node and edge compatibility matrices corresponding to a pair of graphs. Of course, the node and edge compatibilities could potentially be defined in any number of ways, using a variety of information concerning the objects.

**Figure 1:**
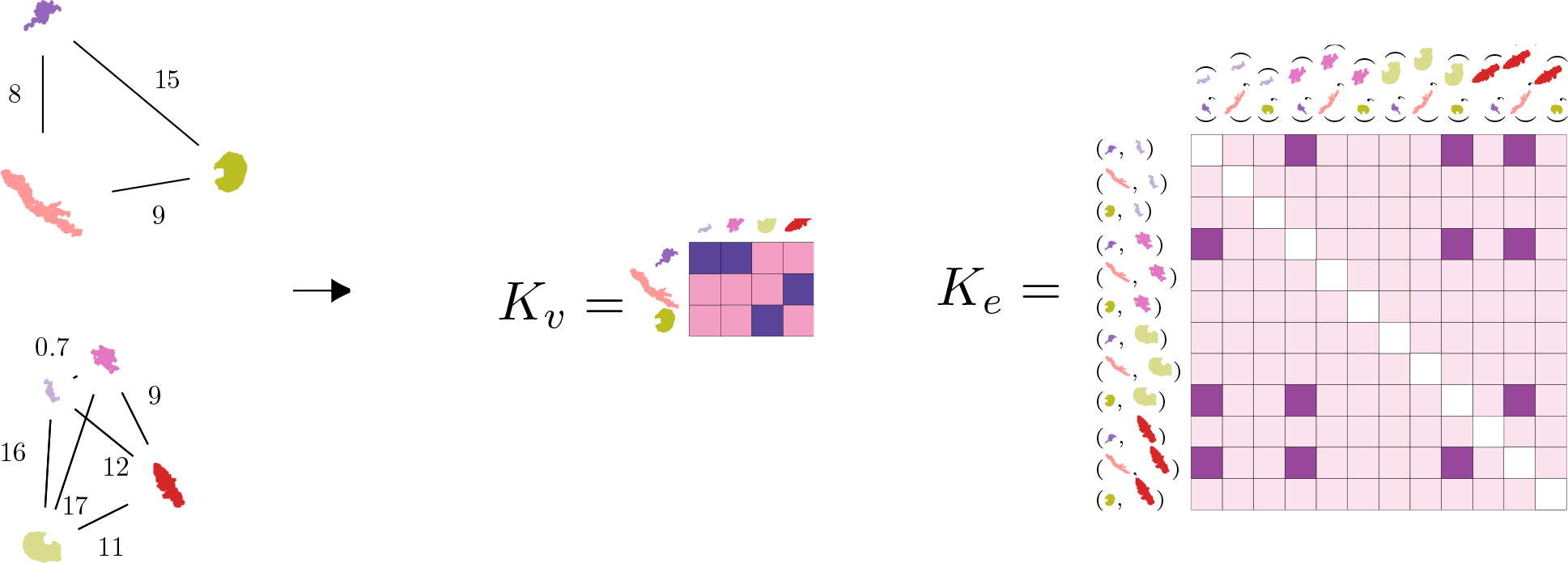
Illustration of the generation of the matrices used in graph matching. The reference frame and a new frame are represented as complete graphs, where the nodes are the objects and the edge weights are the distances between the nodes. The matrix *K*_*v*_ is generated by comparing the similarity of each node in the reference graph (top) with each node in the new graph (bottom). The matrix *K*_*e*_ is generated by comparing the compatibility of each possible match of objects between the two graphs with each other possible match. Darker colors in *K*_*v*_ and *K*_*e*_ indicate larger values.

**Figure 2:**
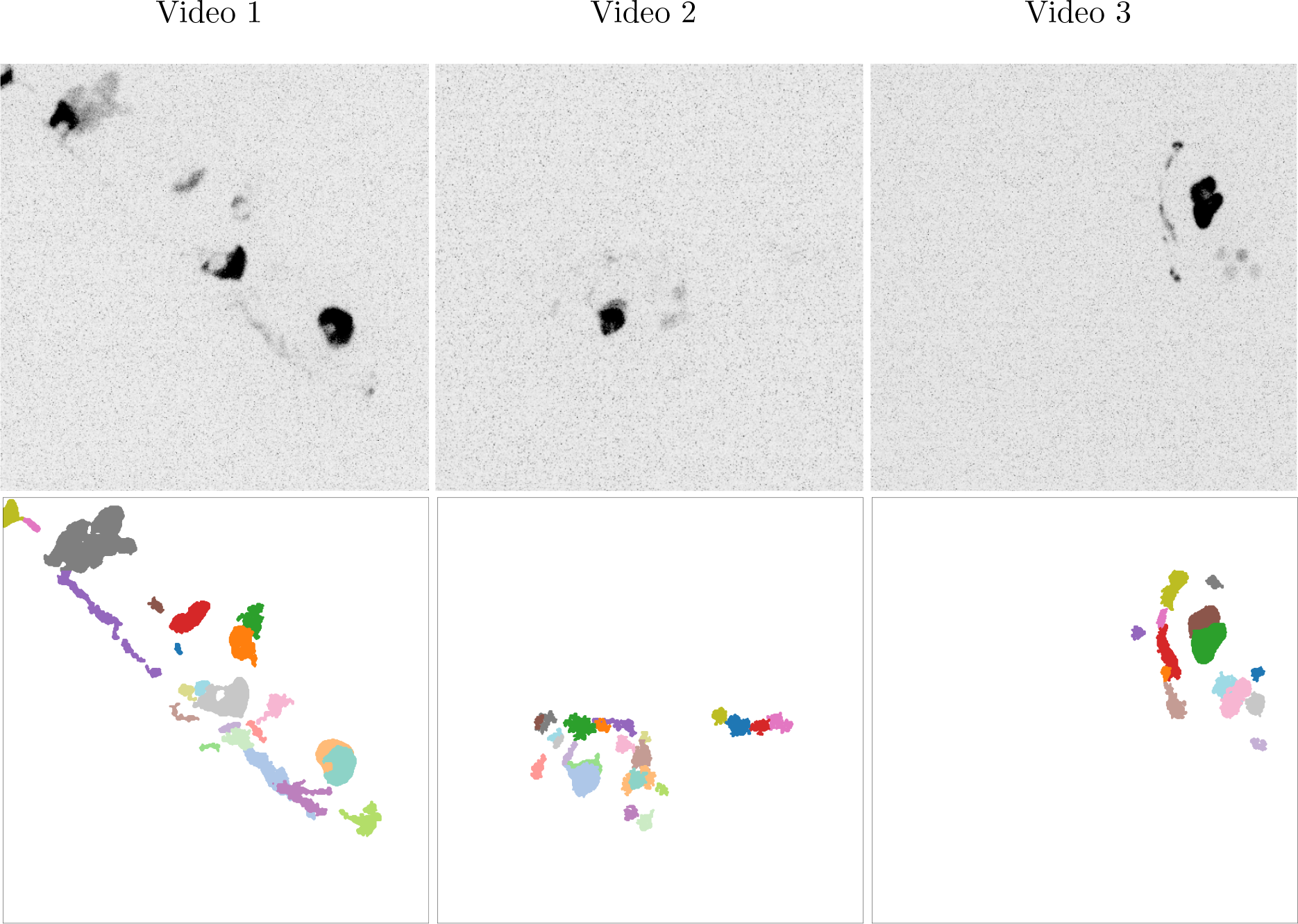
Max-projections of the original reference frames from each video (top row) and the corresponding ground-truth segmentations (bottom row). Note that the colors of individual objects can differ across the videos.

### 2.2 Graph matching formulation

We seek to find an optimal assignment of each node in a given frame to a node in the reference frame using the node and edge compatibility matrices defined above. For such a purpose, two main graph matching formulations exist in the machine learning literature. The first formulation aims to minimize the distance between the adjacency matrix of one graph and a permuted version of the adjacency matrix of a second graph (e.g., Umeyama, 1988). The second formulation, which is more general, aims to maximize the compatibilities of pairs of potential matches (e.g., Gold and Rangarajan, 1996). See Zhou and la Torre (2016) for a discussion of prior work on graph matching. We will build on the second approach, as it is more general, and modify it to produce a novel formulation allowing for many-to-one matching. For now we will assume that the hyperparameters are fixed and will postpone the discussion of how to learn them to Section 2.5.

Denote the number of nodes in the reference frame by *m* and the number of nodes in the given frame by *n*. We will represent a set of possible assignments by a matrix *P* ∈ { 0, 1}^*m*×*n*^. An entry (*i, i*′) in *P* is equal to one if nodes *i* and *i*′ should be matched and is zero otherwise. As each object in the given frame is assumed to occur in the reference frame, each column of *P* should sum to one. We may moreover wish to bound the sum of the rows using a vector *m*_max_ ∈ ℕ^*m*^ indicating the maximum number of allowable matches for each object in the reference frame. When entries in *m*_max_ are larger than one, this allows for an object in the reference frame to have been split into multiple parts in the given frame. This happens fairly often with neurites due to their faintness in the images.

Given the above constraints and the compatibility matrices, we can formulate the problem as a quadratic programming problem:

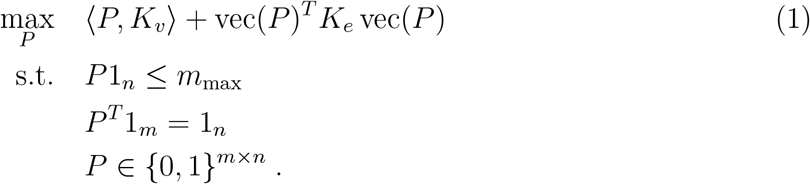

Here vec(*P*) is the vectorization of *P* obtained in the usual column-wise order.

### 2.3 Linear programming relaxation

Problem (1) is NP-hard due to the discrete constraints on *P*. Moreover, due to the edge compatibility term, the objective function is non-convex. Numerous methods have been proposed to solve similar problems, called quadratic assignment problems (Lawler, 1963, Burkard et al., 1998, Loiola et al., 2007). We choose to relax the problem to a linear program by re-writing the quadratic terms in the objective as a function of new variable *Q*, which ideally satisfies *Q* = vec(*P*) vec(*P*)^*T*^, in which case ⟨*Q, K*_*e*_⟩ = vec(*P*)^*T*^ *K*_*e*_ vec(*P*), relaxing the discrete constraints on *P* and *Q* to positivity constraints, and solving the resultant linear optimization problem with respect to the pair *P, Q* (Lawler, 1963, Adams and Sherali, 1986). To obtain a tighter relaxation, we can add several constraints on *Q* based on the constraints on *P* (Frieze and Yadegar, 1983, Torresani et al., 2013). Details regarding our relaxation may be found in Appendix B.1.

### 2.4 Multi-step optimization procedure

In practice, solving the relaxed version of Problem (1) is still intractable for large problems, as it scales quartically with the number of nodes in a graph. Therefore, we instead apply a multi-stage procedure. In the first step, we apply a threshold on the sizes of the objects in each graph and solve the relaxed version of Problem (1) only for the large objects. In addition to reducing the computational complexity, this allows us to focus on the matches for which we should be more confident. This is because the large objects tend to be somas, and somas tend to have more distinct shapes and be segmented as one object instead of multiple objects.

Once we have matched the largest objects, we fix those matches and solve a revised version of Problem (1) on the medium sized objects, which are defined using another threshold. In particular, if we order the previously-assigned objects such that they come first, we can write *P* = [*P*_*m*_, *P*_*u*_], where *P*_*m*_ corresponds to the part of *P* for the already-matched larger objects and *P*_*u*_ corresponds to the part of *P* for the yet-to-be-matched objects. We can similarly write *K*_*v*_ = [*K*_*vm*_, *K*_*vu*_] and *K*_*e*_ = [*K*_*emm*_, *K*_*emu*_; *K*_*eum*_, *K*_*euu*_], where, e.g., *K*_*emu*_ corresponds to the upper right part of *K*_*e*_, which contains terms related to edges in the given frame where one node is matched and the other is not. Since the edge compatibilities that are most likely to be informative are those between unmatched and matched nodes rather than between multiple unmatched nodes, we omit the quadratic term in this phase of the optimization. We furthermore omit the distance constraints. That is, we write the optimization problem corresponding to (1) as

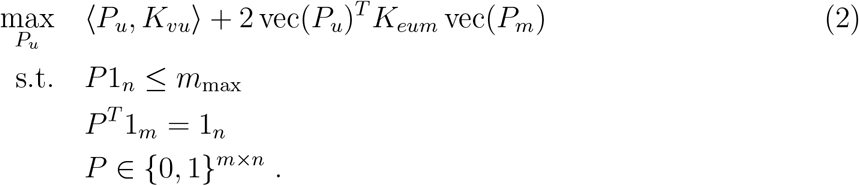

This problem is a linear assignment problem, and we solve it in the same way in which we solve the linear assignment problem detailed in Appendix B.1. Finally, we repeat this step one more time with the remainder of the objects, thereby obtaining matches between all objects in the given frame and objects in the reference frame.

### 2.5 Learning the hyperparameters

Thus far we have assumed that the hyperparameters were fixed. However, we would expect the performance to improve if we have numerous labeled frames and use them to tune the weights of each feature. We do this for each step in the multi-step procedure using ideas from structured output learning. See Appendix B.2 for more details.

The overall matching procedure is outlined in Algorithm 1 in Appendix B, while the learning of the hyperparameters is summarized in Algorithm 2 in Appendix B. As noted in Section 2.4, we perform the learning for three different cutoff sizes of objects. These sizes are determined based on the performance on the training and validation sets when trying a grid of different object sizes. For each pair of cutoff sizes we learn to match the training frame to the reference frame by optimizing the hyperparameters based on Problem (5) in Appendix B.2.

## 3 Implementation

We will now apply our method to three 3D videos of *C. elegans*. Through the numerical experiments, we will demonstrate the feasibility of our method and evaluate the importance of the design choices that were made. In this section we first review the data and experimental setup, and then proceed to discuss the results.

### 3.1 Data

The images in each video were obtained using an inverted spinning disk confocal microscope. The exposure time of each image in a 3D volume was approximately 10 ms Each *C. elegans* worm was a freely-moving young adult hermaphrodite, and a red fluorescent protein was expressed in the nuclei and cytosols of selected interneurons. This is in contrast to much of the literature on tracking *C. elegans*, in which the worms are often (semi-)immobilized, and in which the fluorescence appears only in the nuclei. For more details, see Appendix C.

Figure 2 displays the reference frame taken from each of the original videos (top row), along with the associated segmentations (bottom row). The identities of the segmented objects were manually corrected, and each object is associated with one color. Note that the dark purple object in Video 1 is composed of multiple non-contiguous parts. To deal with this in other frames, our methodology allows for many-to-one matching. Also note that some of the smaller objects, such as the dark brown and dark blue objects in Video 1, will not appear in every frame of the video. Therefore, not every object in the reference frame will be matched each time.

The three videos on which we evaluate the results were chosen to represent a range of possible settings. The number of objects in the reference frame for Videos 1, 2, and 3 is 23, 23, and 15, respectively. In Videos 1 and 3 the worm moves and has a diverse set of postures throughout the video, whereas in Video 2 the worm moves much less. On the other hand, the segmentations are the least consistent for Video 2 due to the overlapping nature of the objects. As we will see in the results section, it is the inconsistency of the segmentations that hurts the performance the most and that makes Video 2 the most difficult to handle. The segmentations are most consistent for Video 3, and this is the video for which we obtain the best results.

### 3.2 Data analysis setup

Prior to running graph matching, preprocessing of the videos was performed using the watershed method to segment the objects; see Appendix C. Moreover, a reference frame was manually chosen for each video, with the criterion that it should contain all or nearly all objects that appear throughout the video. The data was divided sequentially into training, validation, and test sets. The validation set was used to determine the size cutoffs for small, medium, and large objects; see Algorithm 1 in Appendix B. The number of labeled training frames used is indicated in the plots; it varied between 1 and 8. For more details on the setup and the preprocessing, see Appendix C.

We evaluate the performance of the method based on the accuracy of the object labels. Here the segmentation of the objects is taken as given from the preprocessing step. Therefore, it is possible that neurons and neurites could get split into multiple pieces. When computing the accuracy we treat each piece equally.

### 3.3 Results

#### Overall performance

Figure 3 displays the test accuracies for each video when varying the number of labeled training frames between one and eight. The boxplots in green show the results from our overall algorithm, in which the hyperparameters are learned. The average performance with a single labeled frame varies from 68% for Video 2 to 89% for Video 3. With eight labeled frames, these values increase to 76% for Video 2 and 93% for Video 3, respectively, demonstrating the usefulness of having additional labeled frames. Relative to Video 2, Video 3 has fewer total objects and is also more consistently segmented, which lead to the higher accuracies for Video 3. The average runtimes for matching a single frame to the reference frame for fixed hyperparameters are displayed in Table 2 in Appendix D. Across the three videos, matching one pair of frames took 1.4-15.1s on average on a machine with an Intel Xeon E5-2695 v4 processor and 120 GB RAM.

**Figure 3:**
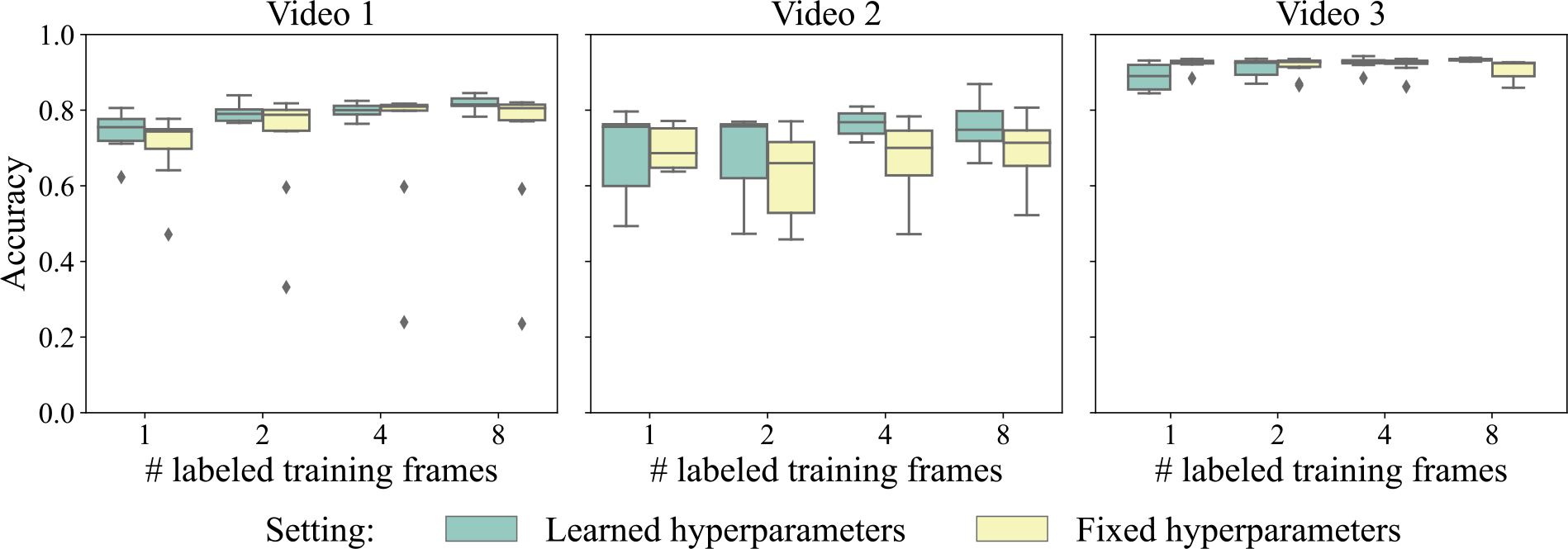
Test set accuracies across ten trials of the proposed algorithm when varying the quantity of labeled training frames and the video the method is tested on. The header above each plot indicates the video the method was tested on.

#### Benefit of hyperparameter learning

There are two factors that can influence the increase in accuracy with the number of labeled frames. First, with more training frames we obtain a better estimate of the maximum number of parts per object (line 10 in Algorithm 2). Second, with more training frames we can better learn the hyperparameters. To examine the impact of each factor, we also include in Figure 3 the performance when fixing the hyperparameters at their initial values. In this setting, with the exception of the maximum number of parts per object, the hyperparameters were fixed at their initial values.

Both of the aforementioned factors contribute to the performance of the algorithm. When tuning only the maximum number of parts per object and fixing the other hyperparameters, the test accuracy changed by -2-3% on average, depending on the video, when going from one to eight labeled frames. In contrast, learning the other hyperparameters resulted in a more varied performance across videos. On Video 1 the accuracy ranges from 5% better on average in the case of one labeled frame to 13% better in the case of eight labeled frames. In contrast, for Video 2 the average accuracy changed by -2-13%. Learning the hyperparameters had the least effect for Video 3, where the performance varied between 4% worse (in the case of a single labeled frame, probably due to overfitting) and 3% better (in the case of 8 labeled frames). These results suggest that the default parameter values worked well for Video 3. Overall, however, in the cases where the initial parameter values are sub-optimal, learning the hyperparameters can significantly improve the performance. We also explore the impact of training on different videos in Appendix D.

#### Benefit of the chosen features

Figure 4 displays the results of an ablation study in which we omit sets of features from graph matching and leave all of the hyperparameters not associated with those features at their previous values. For each video the node features (based on the eccentricity, solidity, and z-level) as a whole play an important role. Without the node features the average accuracy drops by 7-14% for Video 1, 24-32% for Video 2, and 4-10% for Video 3. Among each of these features, the z-level plays the most important role. Without using the z-level at the first iteration of the algorithm, the average accuracy drops by 3-7% for Video 1, 10-15% for Video 2, and 0-3% for Video 3. The edge features (based on the pairwise distances between nodes) also play a very important role in the performance. Without them, the average accuracy drops by 13-25% on Video 1, 8-41% on Video 2, and 8-11% on Video 3. Overall, we find that it is important to include both the node and the edge features in the model.

**Figure 4:**
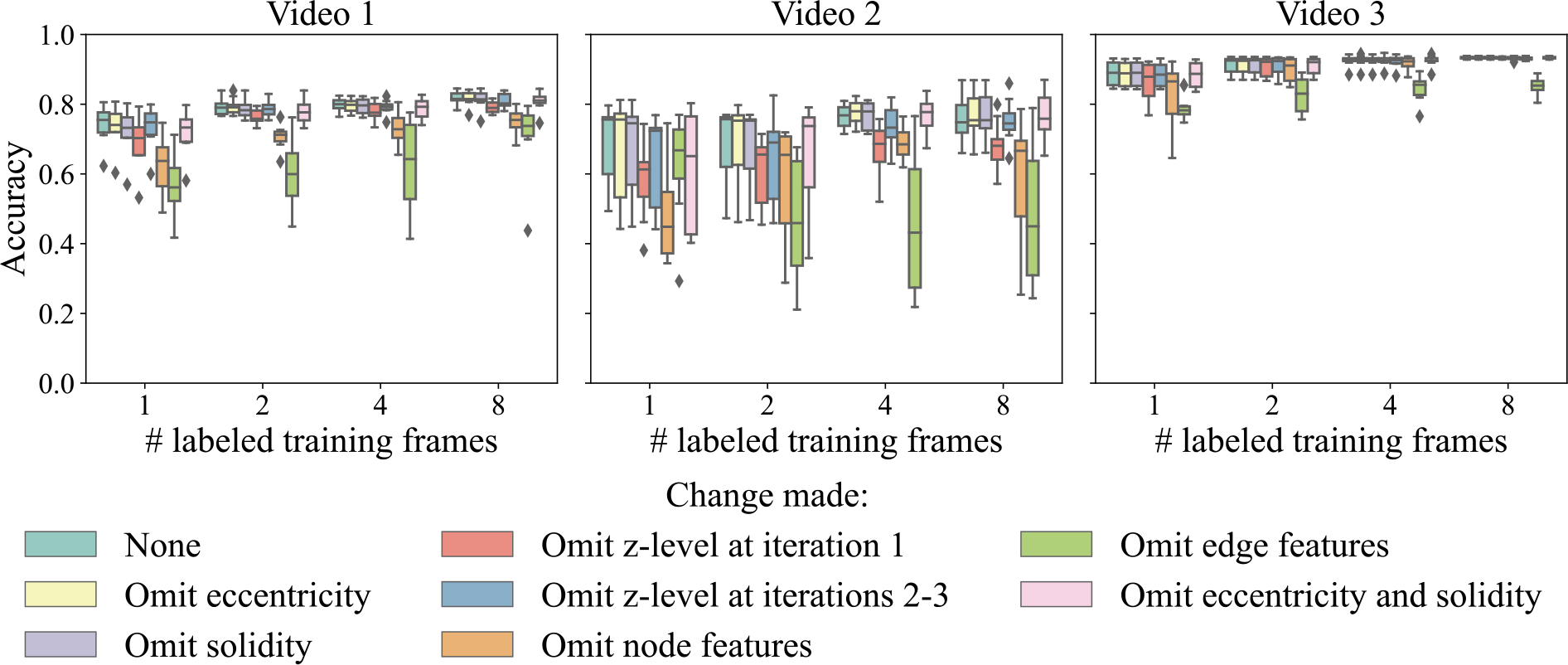
Test set accuracies across ten trials of the proposed algorithm when varying the feature set, the quantity of labeled training frames, and the video the method is tested on.

## 4 Discussion

In this work we presented an approach to tracking neurons in videos of freely-moving *C. elegans*. Neurons in freely-moving worms are generally more difficult to track than those in partially-immobilized worms because the worms deform and move in and out of the field of view, and their neurites are not always visible even if they are within the field of view. To tackle these problems, we represented each frame as a complete graph, and generated node and edge features that were mostly rotation- and translation-invariant. We then performed many-to-one matching between each frame and a given reference frame containing all of the neurons of interest. Overall, we found that when learning the hyperparameters associated with the features we could obtain an average accuracy of 68-89% with just two labeled frames (one reference frame plus one training frame) on the three videos we considered, which increased to 76-93% with seven labeled frames (one reference frame, four training frames, and two validation frames). Therefore, this approach has the potential to drastically decrease the amount of time researchers spend hand-labeling neurons in videos.

### Comparison to prior work

The setting and task that our method targets — that of tracking pre-segmented freely-moving worms with fluorescence in the cytosols — are different from those targeted by other methods in the literature on tracking *C. elegans*. As such, it is not possible to directly compare the performance of our method to the performance reported in other papers.

Methodologically, the works of Chaudhary et al. (2021) and Kainmueller et al. (2014) are most similar to ours. Chaudhary et al. (2021) consider several settings with partially-immobilized worms. Such a setting is easier to tackle because the relative position information of each of the neurons is more informative. When using a data-driven atlas with labeled frames from seven other worms, their accuracy was 73.5% on average when tracking 58-69 cells. This figure is similar to the 76-93% average accuracies we obtained with seven labeled frames. When tracking only 16 cells they were able to reach an average of 98% accuracy. Similarly, Kainmueller et al. (2014) focus on matching nuclei in pre-disentangled, pre-straightened worms. They report an average accuracy of 83% with their automatic method when there is a maximum of 558 nuclei.

Setting-wise, our work is most similar to that of Yu et al. (2021) and Park et al. (2022). These authors track neurons in freely-moving worms using deep learning approaches. Yu et al. (2021) focus on tracking nuclei and report an unsupervised accuracy of 79.1% on a video where each volume contains an average of 69 labeled nuclei. This number is similar to the 68-89% average accuracies we obtained when labeling only the reference frame. However, their task is still rather different from ours, as we track both somas and neurites. In contrast, Park et al. (2022) study the same setting we do, in which we aim to track both somas and neurites. With five labeled frames they achieved an average of approximately 78-88% accuracy with five labeled frames. In contrast, with four labeled frames, we achieved accuracies of 69-91%. However, these values are not directly comparable because the former are reported in terms of the “fraction found”, with an object defined to be found if its IoU with the manual annotation is larger than 33%. In contrast, we count each piece of each segmented object separately when computing the accuracy. It is more difficult to achieve better results with the latter metric, as it is the small pieces of neurites that are hard to match. Another noteworthy aspect is that it takes 8-12 hours to train the model of Park et al. (2022) on a single video with a GPU, whereas our method is faster, parallelizable, and runs on the CPU; see Appendix D for details on the runtimes.

### Future directions

A number of extensions could be interesting to pursue in future work. First, multiple reference frames could be used, and object labels could be chosen based on a majority vote of the matches to the reference frames. Second, one could roughly align the frames based on the current matches, and then perform graph matching again with the addition of features requiring aligned worms, e.g., those from Kainmueller et al. (2014) and Chaudhary et al. (2021), in order to further improve the matching accuracy. Third, one could try to develop a method to enforce consistency between the labels in consecutive frames in a way that would not risk potentially decreasing the overall accuracy. Fourth, one could try to learn the segmentations as well. Right now we take the segmentations as given, so the accuracy of the final results is limited by the quality of the initial segmentations. Finally, we would ultimately like to be able to track the neurons in real time while providing feedback to the worms.

## Appendix

### A Additional Related Work

As noted in Section 1, three main approaches to graph matching have been used in the worm-tracking literature. Here we outline each approach, along with its shortcomings when it comes to tracking somas and neurites in freely-moving worms.

One approach to tracking is via statistical models. Some of these models assume that each frame is a rigid transformation of another one, plus noise (Varol et al., 2020, Nejatbakhsh and Varol, 2021). Unfortunately, such an assumption is unreasonable for freely-moving worms, which often bend. To account for such bends, other models assume that each frame is a non-rigid transformation of another frame obtained via, e.g., thin-plate splines (Nguyen et al., 2017, Wen et al., 2021). However, the method of Nguyen et al. (2017) was not designed to work with objects that can be split into multiple parts. On the other hand, Wen et al. (2021) only make use of the locations of the objects rather than their features. An alternative approach is to model the relative positions of the neurons via a Markov random field (Tokunaga et al., 2014, Hirose et al., 2018). The downside of the method of Tokunaga et al. (2014) is that the only features used to perform the registration are the coordinates of the centroids of the nuclei. In order to apply it to moving worms, the worm would have to first be straightened and aligned. Moreover, work from the graph matching literature, discussed below, demonstrates that adding additional information can improve the performance (Chaudhary et al., 2021). On the other hand, Hirose et al. (2018) assume that objects are not split into multiple parts during the video.

A second approach to tracking is via graph-based approaches. Such methods can be viewed as a subset of statistical model-based approaches, as they can be derived from the notion of conditional random fields (Sutton and McCallum, 2012). They model each image as a complete graph, where the nodes represent nuclei or (parts of) neurites. The goal is to use characteristics of the nodes and edges in a graph to match it with one or more other graphs. Many authors only consider the problem of matching nuclei, and hence make an important assumption that each object can be matched to at most one object in another frame (Lyzinski et al., 2016, Swoboda et al., 2017, 2019). However, when tracking neurites it is unreasonable to assume each neurite will be segmented into the same number of parts each time. While several other authors also make this assumption, it could be relaxed in the context of their approach (Kainmueller et al., 2014, Toyoshima et al., 2020, Chaudhary et al., 2021). Among these approaches, two kinds of features are used: (a) pairwise node features, which indicate the similarities of single nodes in one graph with single nodes in another graph; and (b) pairwise edge features, which indicate the similarities of single edges in one graph with single edges in another graph. Toyoshima et al. (2020) use only the former, whereas Chaudhary et al. (2021) demonstrate the importance of the latter.

A third, more recent, approach to tracking is via deep learning approaches. One main hurdle with deep learning approaches is that they tend to require large amounts of labeled training data, which do not exist in this domain. To overcome this, Yu et al. (2021) and Park et al. (2022) both propose methods for generating artificial labeled images by transforming segmented images to look like other images in the dataset. However, the two methods differ in the structure of the networks and the amount of supervision: Yu et al. (2021) use a transformer model and require no manually-labeled images, whereas Park et al. (2022) use a convolutional network and require some manually-labeled images. The reason why Park et al. (2022) require manually-labeled images is that, unlike Yu et al. (2021), they use the network to perform segmentation in addition to detection. The downside to such deep learning approaches is that the models would need to be trained anew on each strain, or even each worm, a process that can take 8-12 hours on a GPU. In addition, the results are significantly less interpretable than for model-based or graph-based methods. Finally, it is unclear how well the method of Yu et al. (2021) would work in the presence of inconsistent segmentations, such as when one object is sometimes segmented into different numbers of parts. To our knowledge, Park et al. (2022) are the only ones who address the problem of tracking both nuclei and neurites.

### B Algorithmic Details

In this appendix we provide additional details related to the implementation of the overall graph matching algorithm. Algorithm 1 outlines the overall graph matching procedure, while Algorithm 2 describes the method for learning the hyperparameters.

#### B.1 Optimization details

As noted in Section 2.2, Problem (1) is NP-hard because of the constraints on *P*. We therefore proceed by relaxing the problem to a linear program, as described next.

Throughout this derivation, we will assume *Q* = vec(*P*) vec(*P*)^*T*^ and *P* ∈ {0, 1}^*m*×*n*^. First consider the constraint *P*^*T*^ 1_*m*_ = 1_*n*_. A subset *Q*_*·,jm*:(*j*+1)*m*_ of the matrix *Q* for some *j* ∈ {1, …, *n*} satisfies 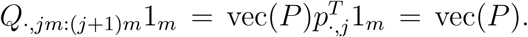. We can derive a similar result when considering *Q*_*im*:(*i*+1)*m,·*_ for some *i* ∈ {1, …, *m*}. We additionally have information regarding *m* × *m* blocks on the diagonal of *Q*. In particular, note that in this case, 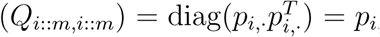_*·*_ and for *i* ≠ *j* 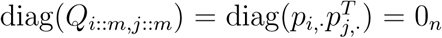, where the notation *i* :: *m* indicates that we take every *m*’th element, beginning at *i*. Finally, we can include the constraint *Q*_*i,j*_ ≥vec(*P*)_*i*_ + vec(*P*)_*j*_ −1.

Including all of the above constraints, we obtain the linear program

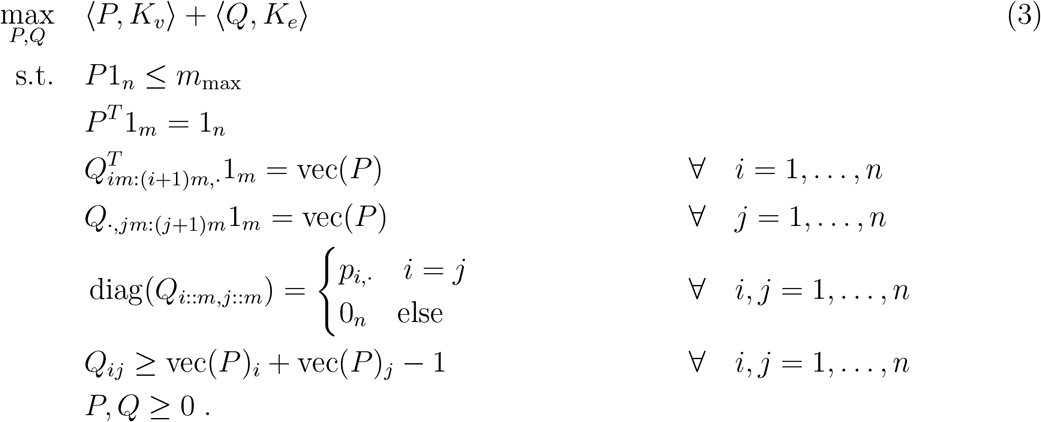

Problem (3) is a linear program and hence one can find its global minimum. In practice we solve it using the version of the high performance dual revised simplex method implemented in Scipy (Huangfu and Hall, 2018, Virtanen et al., 2020). However, the minimizer of Problem (3) will not in general be a feasible solution to Problem (1). Therefore, we round the solution by solving another linear program. Specifically, letting 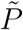 be the solution to Problem 3, we then solve the problem

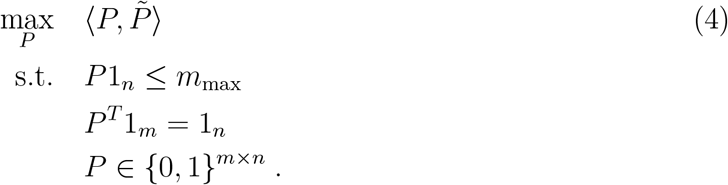

This can be viewed as a linear assignment problem. We solve it using a modified version of the Jonker-Volgenant algorithm that is implemented in Scipy (Crouse, 2016, Virtanen et al., 2020).

#### B.2 Learning the hyperparameters

Numerous researchers, including Taskar et al. (2005), Tsochantaridis et al. (2004) and Pillutla et al. (2018), have introduced frameworks for learning hyperparameters in structured prediction problems such as graph matching via the use of a structural support vector machine. We apply these ideas to learn hyperparameters in our setting.

Denote the set of hyperparameters by *w*, the *i*th (labeled frame, reference frame) pair by *F* ^(*i*)^, and the corresponding matches by *P* ^(*i*)^, *i* = 1, …, *L*. Furthermore, let *f* (*F* ^(*i*)^, *P* ; *w*) denote the objective value from either Problem (3) or Problem (2) (depending on the stage of the procedure) when the matches are given by *P* and the hyperparameters are *w*. Finally, denote the space of all possible matches for the *i*th frame by 𝒫^(*i*)^ and let *ℓ*(*P* ^(*i*)^, *P*) = *n* −⟨*P* ^(*i*)^, *P* ⟩be the number of incorrectly-labeled objects in frame *i* based on *P* when the true matches are given by *P* ^(*i*)^. A general form of the hyperparameter learning problem takes the form

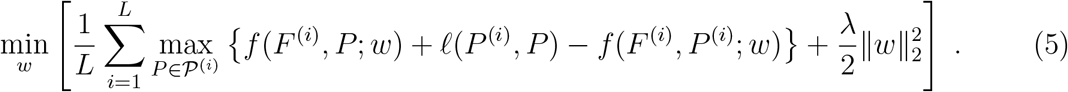

Roughly speaking, for fixed *w* the method attempts to find the matching *P* that most grossly dissatisfies the inequality *f* (*F* ^(*i*)^, *P* ^(*i*)^; *w*) ≥ *f* (*F* ^(*i*)^, *P* ; *w*) + *ℓ*(*P* ^(*i*)^, *P*). This constraint says that *P* ^(*i*)^ should always be better than any other potential matching *P* (based on the original objective function *f*), and that there should be a margin *ℓ*(*P* ^(*i*)^, *P*) between the function values at *P* ^(*i*)^ and any other *P*. Now, for fixed *P*, the problem attempts to make the original objective function *f* evaluated at *P* as small as possible relative to *f* evaluated at the true *P* ^(*i*)^, subject to a penalty that prevents the hyperparameters from becoming too large. In practice we perform alternating minimization to find a local optimum of Problem (5), optimizing over *P* for fixed *w* via a linear programming solver and then optimizing over *w* for fixed *P* using SVRG, a variance-reduced version of stochastic gradient descent (Johnson and Zhang, 2013).

##### Algorithm 1

Full Matching Procedure

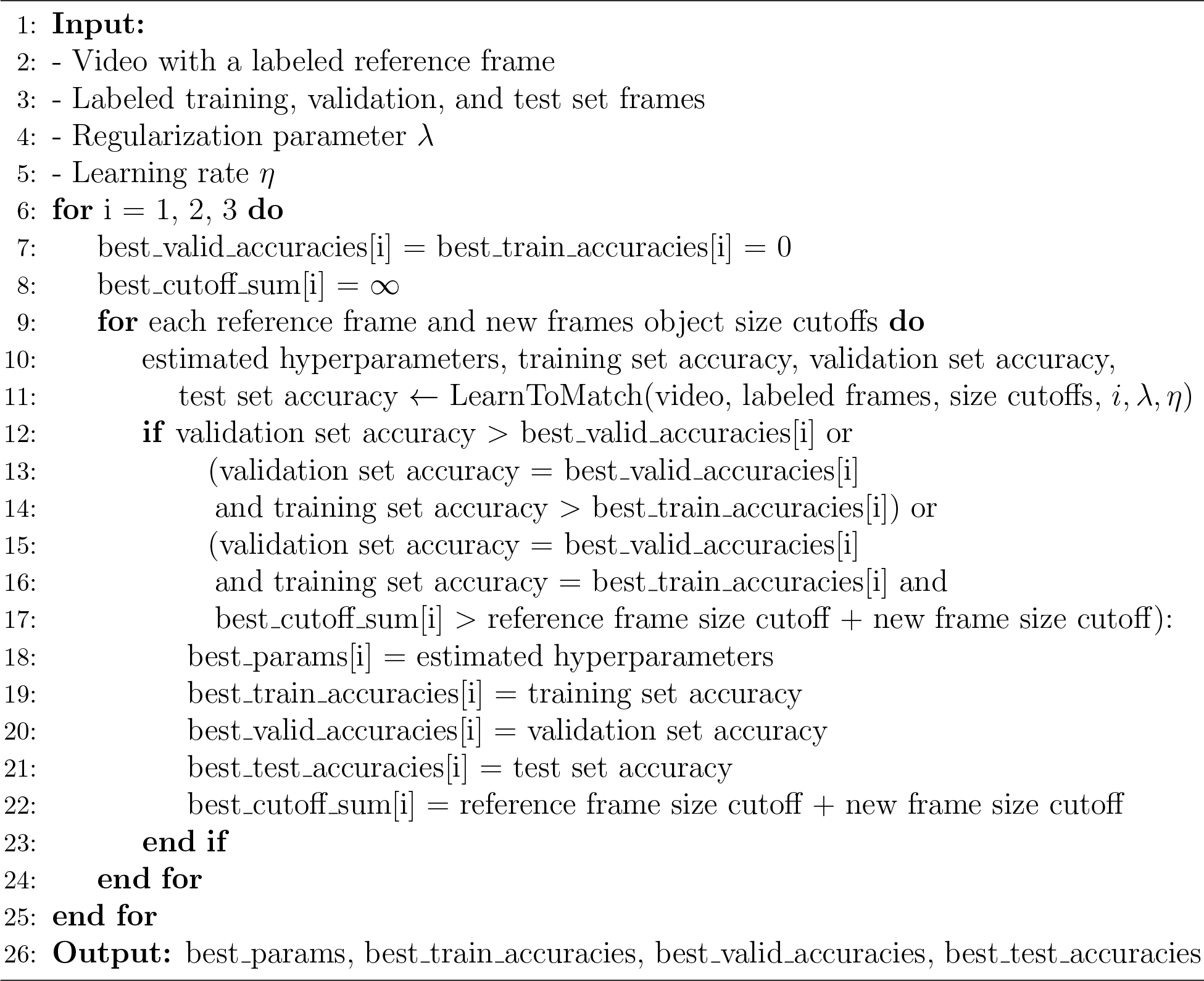

### C Additional Experimental Details

In this appendix we provide additional details related to the data used and the setup of the numerical experiments.

#### Data

We performed calcium imaging of *C. elegans* hermaphrodites as detailed in Park et al., 2022. In the recordings, from which snapshots are shown in Figure 2, we used worms of strains SJR16, marking the second layer interneurons, and SJR15, marking the first and second layer interneurons. The genotypes of each strain were as follows: SJR16: *sjxIs9[* P*npr-9::NLS-wrmScarlet;* P*ttx-3::mNeptune::GCaMP6s;* P*ceh-16::mNeptune::GCaMP6s;* P*gcy-28d::mCherry::GCaMP6s;lin-15]* ; *aeaIs009[* P*rgef-1:NLS-GCaMP6s]*. SJR15: *sjxIs8[* P*npr-9::NLS-wrmScarlet;* P*ttx-3::mNeptune::GCaMP6s;* P*ceh-16::mNeptune::GCaMP6s;* P*gcy-28d::mCherry::GCaMP6s;lin-15]; aeaIs009; sjxIs9*.

#### Preprocessing

Prior to performing graph matching, the red channel from each video was preprocessed following a similar approach to the one taken by Park et al. (2022). See Table 1

**Table 1:** Summary of the parameter values used during the data preprocessing, in the order in which they are used.

| Parameter | Video 1 | Video 2 | Video 3 |
| --- | --- | --- | --- |
| Rows removed | {511, 512} | {434} | {514} |
| Columns removed | {} | {433, 434} | {515, 516} |
| $\sigma_1$ in the difference of Gaussians | 1.5 | 1 | 1 |
| $\sigma_2$ in the difference of Gaussians | 15 | 25 | 25 |
| Thresholding quantile | 0.97 | 0.97 | 0.95 |
| Max object volume (pixels) | 100000 | 100000 | 100000 |
| Min object volume (pixels) | 120 | 120 | 120 |
| Min volume of a large object (pixels) | 500 | 500 | 1200 |
| Max distance to a large object (pixels) | 10 | 10 | 12 |
| $\sigma$ in the Gaussian filter | 5 | 5 | 5 |
| Threshold for merging objects | 0.2 | 0.2 | 0.2 |
| Min final object volume (cubic microns) | 3 | 3 | 3 |

##### Algorithm 2

Learn To Match

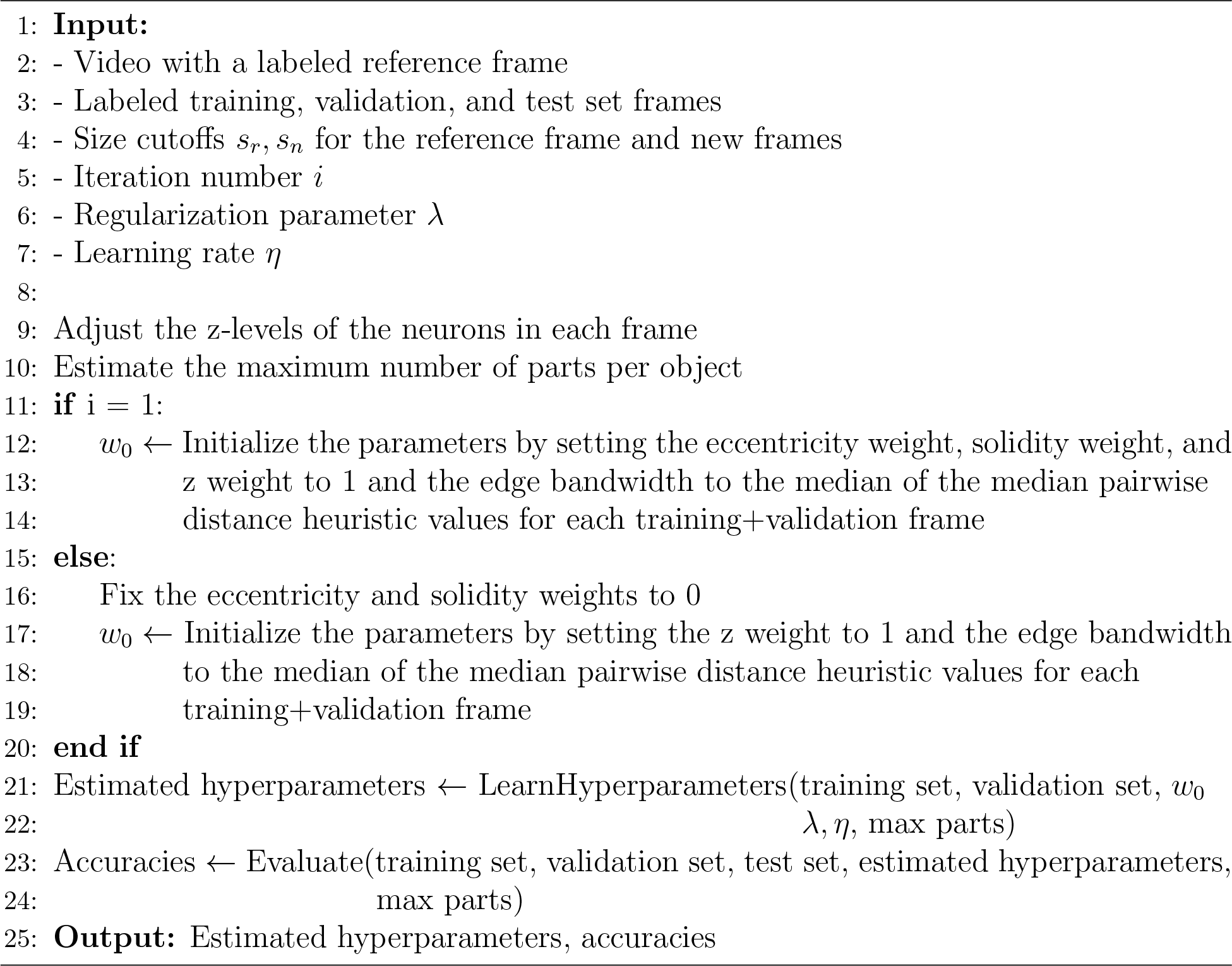

for the values of the parameters used. We first remove bad fibers (the tensor equivalent of rows and columns) from the images. These are identified visually via 2D max projections of the images, and in them all pixels are typically zero. Next, the images are smoothed using a difference of Gaussians filter and then thresholded such that only pixels above a specific quantile are retained. To remove noise, we keep only the connected components that are larger than a given lower bound (120 pixels) and smaller than a given upper bound (100000 pixels). In a further effort to eliminate noise, we then remove connected components far (10-12 pixels) from any “large” object in the image (defined to be either 500 or 1200 pixels in size, depending on the video).

Once the image is thresholded and the noise is removed, we apply the watershed method to identify the objects in the images (Forsyth and Ponce, 2012). The distance transform is applied using the relative resolutions of the axes (0.1625×0.1625× 1.5), and then a Gaussian filter is applied to the result. The local maxima are identified within local regions of size 13×13×3. As the watershed method tends to oversegment the images, we postprocess the results. In particular, we merge neighboring objects if the number of touching pixels divided by the approximated surface area (given by 2/3 of the volume in pixels) is larger than a given threshold. Finally, we remove objects smaller than a minimum acceptable volume (in microns).

##### Extraction of relevant quantities

Following the aforementioned preprocessing, we extract the quantities from each frame that will be used in the graph matching and the post-processing of the matches. We compute the center of mass along each axis, scaling the resultant values based on the resolution of each axis in the images. Pairwise distances between objects are computed based on the objects’ centers of mass. We then compute the solidity and eccentricity of each object using scikit-image’s regionprops function (van der Walt et al., 2014). Finally, we compute the volume of each object based on the number of pixels it contains.

The algorithm assumes that the centers of mass in the z-direction are roughly constant over time. However, in reality this is not the case due to (1) the up-and-down motion of the microscope stage; and (2) the motion of the worm. To adjust for (1), we difference the median center of mass in the z-direction of the large objects in the odd frames and that in the even frames. The difference is then added to all z-values in the even frames. To adjust for (2), we then smooth the median centers of mass in the z-direction across frames and subtract the smoothed value for each frame from the z-values. The smoothing is performed using a median filter of size 9.

##### Reference frame

A single reference frame was chosen for each video. Each such frame was chosen by hand and was such that it contained nearly all of the objects seen throughout the video. The segmentation of the reference frame was manually corrected. If there were several objects that touched each other and that would be impossible to consistently segment, these objects were merged into one. It is possible that such objects could be separated during a post-processing stage. Objects that were missing completely from the reference frame were not artificially added.

##### Creation of the training/validation/test sets

In each of the three datasets used, we manually annotated the object identities in 150 frames (but did not correct the segmentations). Due to the high autocorrelation between frames, we elected to use every 5th frame, beginning with the first frame and ending with approximately the 746th frame (frames containing no objects were removed prior to the frame selection). Unless otherwise specified, the training frames were chosen at random from the first 64 of the 150 annotated frames. The validation frames were then chosen at random from the 67th through the 98th annotated frames. The test frames were the last 50 annotated frames. By selecting the frames in this manner, we generated a buffer of 2 annotated frames (10 original frames) between the training and validation sets and between the validation and test sets. The number of validation frames used in each experiment was set to half the number of training frames (rounded down). In each case, there were 50 test frames.

##### Fixed parameters

At each stage of the optimization we only consider objects whose sizes are above a certain threshold. At each stage we have one threshold for the reference frame and another threshold for the remaining frames. At the first stage we consider all possible cutoffs for the reference frame such that there are at least 5 and at most 20 objects. For the other frames we consider a grid with increments of 100 pixels. The minimum and maximum values are chosen such that 95% of frames have at least three objects and at most ten objects. Due to memory constraints, we restrict further restrict the set of possible cutoff pairs to be such that the largest product between the number of objects in the reference frame above the given reference frame cutoff size and the number of objects in a new frame above the given new frame cutoff size to be smaller than 90. Furthermore, to speed up the training, we restrict the reference frame cutoff to be smaller than the cutoff for the other frames. The motivation behind this is that we want to ensure that all objects in the new frame that are above the cutoff are also in the reference frame. For the second stage of the optimization we consider the same grids, but require both the cutoffs to be smaller than the cutoffs in the first stage, and we again require the cutoff for the reference frame to be smaller than the cutoff for the other frames. Finally, for the third stage of the optimization we fix both cutoffs to zero.

In Problem (3) we introduced a constraint on the maximum number of matches per object. We tune these values for each iteration based on the training set and the reference frame. For the first stage of the optimization, we generally use the maximum number of times each object above the given cutoff size is observed in the training frame(s). However, for objects that appear in the reference frame that are not in the training frames, we assign the maximum number of times they can appear to 1. For the second and third stages, we set the cutoffs similarly. However, we add an adjustment value of two for each object, since we some objects may have many parts and we may not observe all of those parts in the training frame(s).

The parameters related to the optimization are as follows. The *ℓ*_2_ regularization parameter *λ* is set to 0.01*/L*, where *L* is the number of training frames. The learning rates at each stage of the optimization are 2^*−*5^*/*(0.25 + *λ*), 2^*−*4^*/*(0.25 + *λ*), and 2^*−*4^*/*(0.25 + *λ*), respectively. During the optimization of the hyperparameters we perform a transformation *w*↦exp(*w*) of the variables *w* in order avoid having to include non-negativity constraints. In the first stage of the optimization we initialize the weights of the eccentricity term, solidity term, and z-level term to 1 (on the original scale). We initialize the bandwidth of the edge features using the median of the median pairwise distance heuristics obtained from the distances between each pair of objects in each new frame in the training set (above the given threshold) and each pair of objects in the reference frame (above the given threshold). We optimize the objective (5) using SVRG for 10 epochs. In the second and third stages of the optimization we fix the weights of the eccentricity term and solidity term to zero, as the eccentricity and solidity tend to be less meaningful for small objects. We initialize the remaining parameters in the same way as before. As the optimization at these stages is faster, we use SVRG for 100 epochs.

##### Learned parameters

The parameters that are learned in Algorithm 1 are the cutoff values of the object sizes at each iteration, along with the eccentricity weight, solidity weight, z-weight, and edge features bandwidth at the first iteration, and the z-weight and edge features bandwidth at the second and third iterations. The cutoff values for the object sizes are chosen via a grid search. The other parameters are learned, each starting from a default value of 1. When performing a grid search to determine which object cutoff sizes at each layer are best, we consider, in order, the validation accuracy, the training accuracy, and the sum of the cutoff sizes (the lower the better).

##### Code

The main code for this project was written in Python using the common packages numpy, matplotlib, pandas, scikit-image, scikit-learn, scipy, and seaborn (Harris et al., 2020, Hunter, 2007, Reback et al., 2021, van der Walt et al., 2014, Pedregosa et al., 2011, Virtanen et al., 2020, Waskom, 2021). The optimization of the hyperparameters was performed using a slightly-modified version of the Casimir package of Pillutla et al. (2018). The numerical experiments were performed on a computing cluster where the nodes had 120 GB RAM and Intel Xeon E5-2695 v4 processors that run at 2.10 GHz.

### D Additional Results

#### Learning from other videos

Labeling frames in new videos can be tedious, so being able to train on other, previously-labeled videos could be useful. In terms of the corresponding results, we may expect there to be a benefit to doing this in the cases where there was a benefit in learning the hyperparameters. We explore this in Figure 5. In this figure we also include the results when fixing the hyperparameters at their initial values. Overall, 10/360 of the experiments failed because the method ran out of memory. As this was due to the cutoffs chosen on the other video, we increased the size cutoff for reference objects until one was found for which the algorithm ran successfully. In each experiment we fixed the maximum number of parts per object to the values obtained when training on the same video. This allows us to separate the impact of estimating the maximum number of parts per object from the impact of estimating the other hyperparameters.

**Figure 5:**
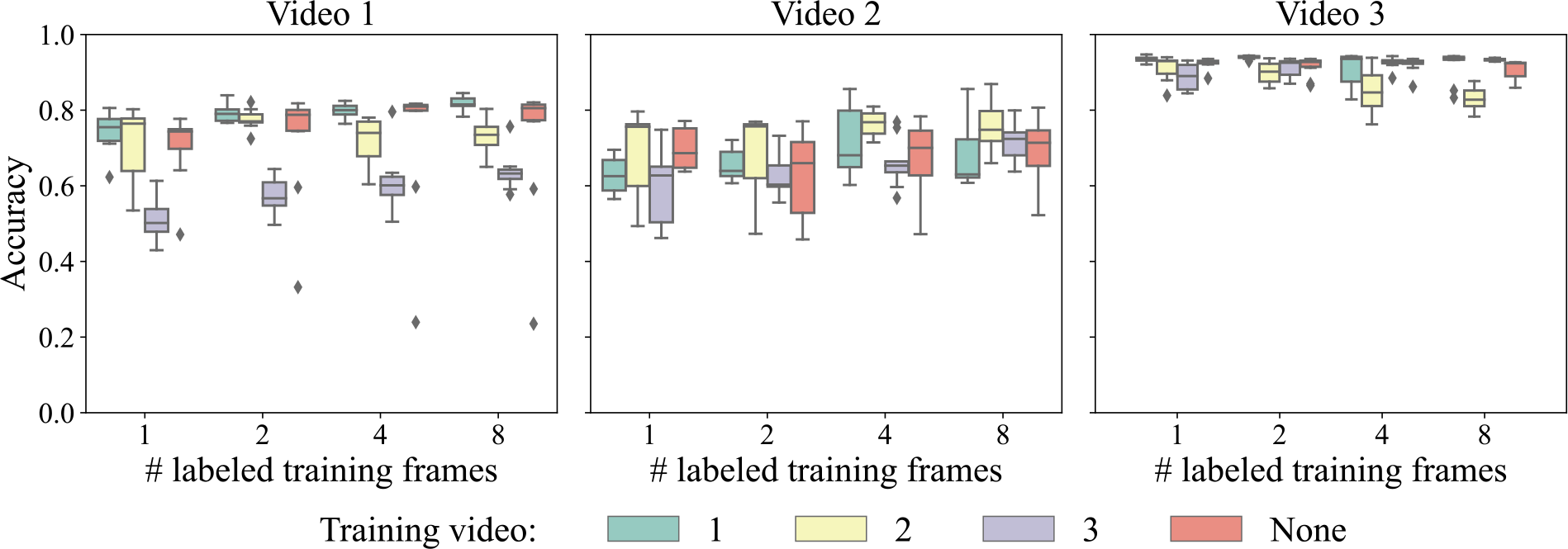
Test set accuracies across ten trials of the proposed algorithm when varying the video used to learn the hyperparameters, the quantity of labeled training frames, and the video the method is tested on. The header above each plot indicates the video the method was tested on.

Figure 5 indicates that training on other videos can be useful for Video 2. In particular, training on Video 1 improved the performance by 4% and 6% relative to not learning the hyperparameters when there were 4 and 8 labeled frames, respectively. In contrast, training on Video 3 with the same numbers of labeled frames led to a 0% improvement and a -2% drop in performance, respectively, relative to not learning the hyperparameters.

#### Runtimes

Generating the features for each frame given the segmentations takes 6-12s per frame, depending on the video, and is parallelizable across all frames. Table 2 summarizes the runtimes of graph matching when the hyperparameters are fixed. The column “Avg. # objects in new frame” indicates the average number of objects (across four different numbers of training frames, 10 random seeds, and 50 test frames) in each new frame being matched to the reference frames. The column “Avg. # objects in ref. frame” indicates the same statistic for the reference frames. As the first iteration entails solving the large linear program given by Problem (3), this iteration takes the longest, at 1-15s per frame, on average. The second and third iterations entail solving the linear assignment problem (2), which is significantly faster. As each frame is independently matched to a reference frame, graph matching for an entire video is embarrassingly parallelizable.

**Table 2:** Summary of the time required to perform graph matching on a single pair of frames for fixed hyperparameters.

| Iteration | Video | Avg. # objects in<br>new frame | Avg. # objects in<br>ref. frame | Avg. runtime (s) |
| --- | --- | --- | --- | --- |
| 1 | 1 | 8.1 | 13.0 | 15.1 |
|  | 2 | 2.7 | 19.1 | 1.4 |
|  | 3 | 8.8 | 10.1 | 7.4 |
| 2 | 1 | 10.5 | 18.5 | $8.6 \times 10^{-4}$ |
| | 2 | 5.9 | 23.0 | $4.8 \times 10^{-4}$ |
| | 3 | 10.0 | 13.6 | $3.0 \times 10^{-4}$ |
| 3 | 1 | 24.0 | 23.0 | $1.6 \times 10^{-3}$ |
| | 2 | 22.0 | 23.0 | $1.0 \times 10^{-3}$ |
| | 3 | 13.5 | 14.0 | $3.9 \times 10^{-4}$ |

Table 3 summarizes the number of (reference frame object size cutoff, new frame object size cutoff) pairs considered at each iteration, and the associated average ranges for the number of objects in the new frames and the number of objects in the reference frames. In the full procedure outlined in Algorithm 1, the hyperparameters are learned, on average, for 120-163 different cutoff configurations at the first iteration and 9-72 cutoff configurations in the second iteration. At the third iteration the size cutoffs are set to zero, so only one configuration is considered.

**Table 3:** Summary of the average number of size cutoff pairs considered for each video at each iteration, along with the associated ranges of average minimum to average maximum numbers of objects in the new and reference frames.

| Iteration | Video | # cutoffs | # new frame objects | # ref. frame objects |
| --- | --- | --- | --- | --- |
| 1 | 1 | 163 | 1-11 | 4-19 |
|  | 2 | 123 | 2-9 | 4-19 |
|  | 3 | 120 | 2-11 | 4-13 |
| 2 | 1 | 72 | 4-17 | 10-22 |
|  | 2 | 49 | 2-15 | 10-22 |
|  | 3 | 9 | 7-12 | 9-13 |
| 3 | 1 | 1 | 12-29 | 23-23 |
|  | 2 | 1 | 17-27 | 23-23 |
|  | 3 | 1 | 12-17 | 14-14 |

## Footnotes

1 We could also have multiple reference frames and perform a voting to determine object identities, at the expense of additional labeling time and computational cost. Such an approach was followed by Nguyen et al. (2017) with non-manually-corrected reference frames.

